# The Hypoxia Response Pathway Promotes PEP Carboxykinase Expression And Gluconeogenesis

**DOI:** 10.1101/2021.05.04.442650

**Authors:** Mehul Vora, Stephanie M. Pyonteck, Tarmie L. Matlack, Aparna Prashar, Nanci S. Kane, Premal Shah, Christopher Rongo

## Abstract

Actively dividing cells, including some cancers, rely on aerobic glycolysis rather than oxidative phosphorylation to generate energy, a phenomenon termed “the Warburg effect^1^.” Constitutive activation of the Hypoxia Inducible Factor (HIF-1), a transcription factor known for mediating an adaptive response to oxygen deprivation (hypoxia), is a hallmark of the Warburg effect^2^. HIF-1 is thought to promote glycolysis and suppress oxidative phosphorylation. Here, we show instead that HIF-1 can promote gluconeogenesis. Using a multiomics approach, we determined the genomic, transcriptomic, and metabolomic landscapes regulated by constitutively active HIF-1 in *C. elegans*. We performed RNA-seq and ChIP-seq under aerobic conditions in mutants lacking EGL-9, a key negative regulator of HIF-1, and then integrated these approaches to identify over a hundred genes directly and functionally upregulated by HIF-1. We show that HIF-1 directly promotes the expression of PCK-1, a PEP carboxykinase that is a rate-limiting mediator of gluconeogenesis^3^. This activation of PCK-1 by HIF-1 promotes survival in response to both oxidative and hypoxic stress. Our work is the first to identify functional direct targets of HIF-1 *in vivo*, and it describes the first complete metabolome induced by constitutive HIF-1 activation in any organism.

## MAIN TEXT

Metazoans respond to hypoxia using a conserved hypoxia response pathway that mediates an adaptive response. Oxygen is sensed by a prolyl hydroxylase (EGL-9 in *C. elegans*), which negatively regulates Hypoxia Inducible Factor (HIF-1 in *C. elegans*), the pathway transcriptional effector (Fig. 1a). When hypoxia ensues, EGL-9 is inhibited, and HIF-1 becomes active. In disorders involving acute hypoxia, tissues are damaged by energy deprivation and oxidative stress^2,4^. The hypoxia response pathway, when activated by hypoxia, minimizes such damage; however, long-term adaptations and tissue remodeling triggered by the hypoxia response pathway itself can also cause damage^5,6^. The pathway can also be activated to promote anaerobic glycolysis (i.e., the Warburg effect) even under aerobic conditions, including in some cancers, in dividing stem cells, in the proliferation of activated T lymphocytes, during endometrial decidualization, and presumably in patients receiving prolyl hydroxylase inhibitors to treat anemia associated with chronic kidney disease^1,7-17^. HIF-1 clearly upregulates enzymes that mediate anaerobic glycolysis, but our understanding of the complete adaptive response mobilized by pathway activation has largely focused on gene expression changes in cultured cells^18-21^. We therefore set our goal on a complete characterization of the adaptive response induced by pathway activation in a whole organism context by taking advantage of the genetic model system *C. elegans*. These nematodes employ single orthologs of the conserved hypoxia response pathway, with null mutants in *hif-1* and *egl-9* being viable but showing altered sensitivity to hypoxic stress^22-29^. HIF-1 is constitutively active in *egl-9* mutants, which we have used here in a combined genomic, transcriptomic, and metabolomic approach to obtain a holistic and functional description of HIF-1 activation in an intact animal.

**Figure 1.**
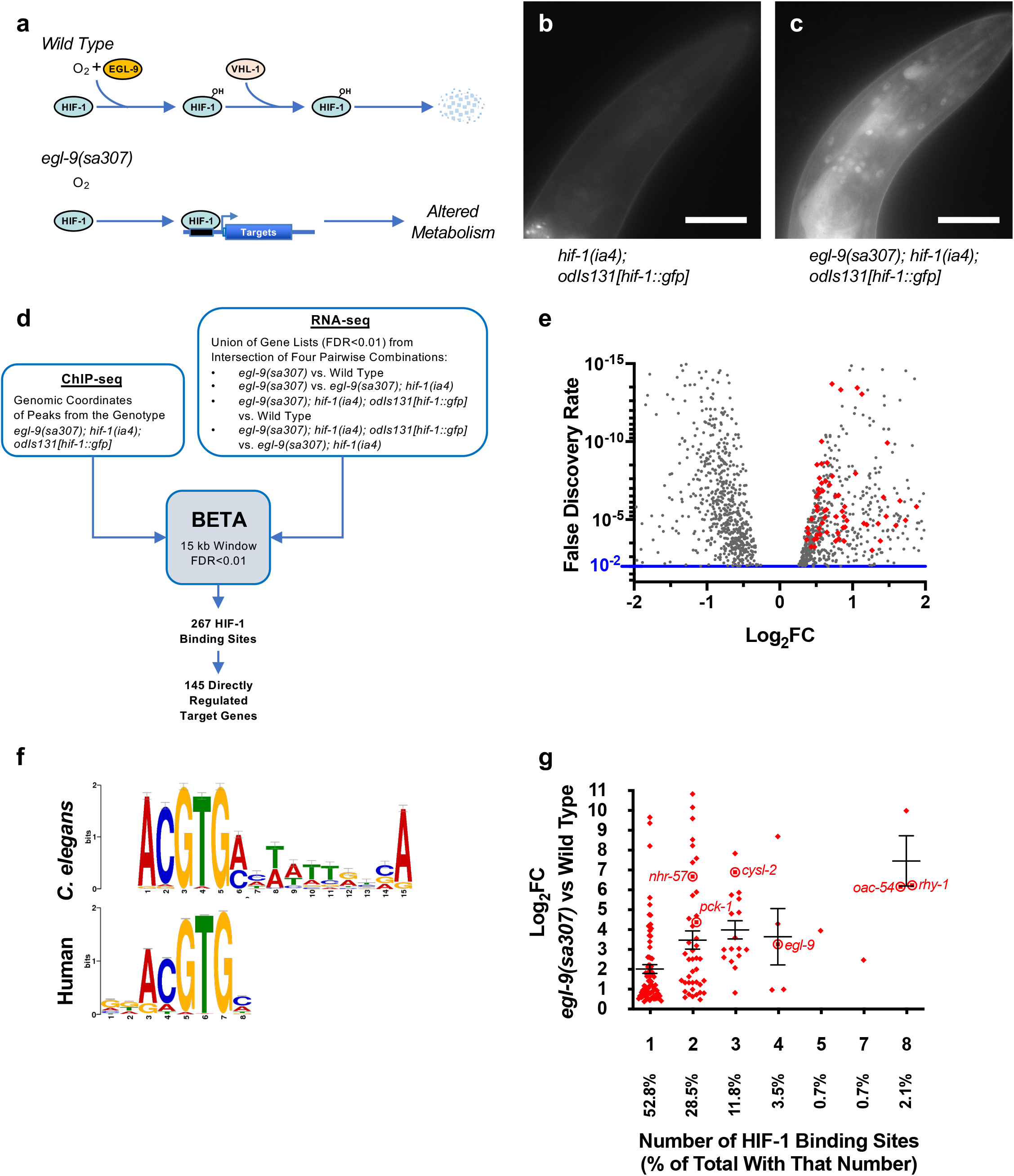
Identifying Genes Directly Regulated By HIF-1. (a) Diagram of the hypoxia response pathway in *C. elegans*. HIF-1 is hydroxylated by EGL-9, ubiquitinated by VHL-1, and degraded by the proteasome. In *egl-9* mutants, HIF-1 remains stable and regulates the transcription of target genes whose expression alters metabolism. (b,c) HIF-1::GFP fluorescence in the indicated genotypes. Bar indicates 30 microns. (d) Strategy for integrating HIF-1 ChIP-seq and RNA-seq data to identify directly regulated targets. (e) Volcano plot of RNA-seq FDR values versus log_2_ fold change expression for individual genes (dots) in *egl-9(sa307)* mutants relative to wild type. The 145 direct targets identified by BETA are indicated with red diamonds. (f) Consensus HRE sequences identified by MEME-suite in humans and enriched in the *C. elegans* ChIP-seq sequences identified by BETA. (g) Graph of log_2_ fold change in expression for individual genes (dots) in *egl-9(sa307)* mutants relative to wild type versus number of HIF-1 binding sites for those genes identified by ChIP-seq. Target genes mentioned in the text are circled and labeled. Error bars indicate SEM.

### A Functional HIF-1::GFP Transgene

To identify sites in the genome bound by HIF-1, we introduced a genomic clone^30^ containing a *hif-1::gfp* chimeric gene expressed from the *hif-1* promoter into the germline genome of *hif-1(ia4)* null mutants (Extended Data Fig. 1a,b). Quantitative mRNA measurements of this genome-integrated *odIs131[hif-1::gfp]* transgene indicated that it is expressed at levels similar to that of endogenous *hif-1* (Extended Data Fig. 1c-e). Under aerobic conditions, HIF-1 is rapidly turned over through the action of EGL-9, which uses oxygen to hydroxylate key proline residues on HIF-1. VHL-1 then ubiquitinates hydroxylated HIF-1, targeting it for degradation by the proteasome^23^. By contrast, HIF-1 is stable in *egl-9(sa307)* null mutants. We observed little detectable fluorescence from *hif-1(ia4); odIs131[hif-1::gfp]* animals under aerobic conditions (Fig. 1b) and nuclear HIF-1::GFP fluorescence in essentially all tissues in *egl-9* mutants (Fig. 1c). The *odIs131[hif-1::gfp]* transgene accurately substituted for endogenous *hif-1* with regard to the regulation of two established transcriptional targets (Extended Data Fig. 1f,g), egg laying (Extended Data Fig. 1h), and survival under hypoxic stress (Extended Data Fig. 1i)^25,26,31-33^. Taken together, these results show that *odIs131[hif-1::gfp]* is functional and expresses tagged *hif-1* to levels similar to that of endogenous *hif-1*, providing a tool to identify genomic binding sites under physiological conditions.

### Identifying Direct HIF-1 Targets

To identify genes directly regulated by HIF-1, we first performed ChIP-seq on L4-stage *egl-9(sa307); hif-1(ia4); odIs131[hif-1::gfp]* animals, which have activated HIF-1 under aerobic conditions, using anti-GFP antibodies to precipitate DNA bound by HIF-1::GFP followed by high-throughput sequencing to identify peaks of reads corresponding to HIF-1 binding sites. We identified 604 peaks (Supplemental Table 1), nearly all of which fell within 500 bps of the transcriptional start site (TSS) or stop codon of a known gene (Extended Data Fig. 2a,b), as well as regions enriched for the hypoxia response element (HRE) known to bind to human HIF1^34,35^ (Extended Data Fig. 2c).

The nearest neighboring gene to a ChIP-seq peak is often not the direct target regulated by the transcription factor bound at that peak. We therefore performed RNA-seq to analyze differential gene expression due to HIF-1 activation, using these data to identify HIF-1-regulated genes near HIF-1 binding sites. We examined transcriptomes of L4-stage animals under aerobic conditions in genotypes in which HIF-1 is active, including *egl-9(sa307)* mutants and *egl-9(sa307); hif-1(ia4); odIs131[hif-1::gfp]* mutants, as well as inactive, including wild type and *egl-9(sa307); hif-1(ia4)* double mutants (Extended Data Fig. 3a). Multidimensional Scaling analysis showed tight clustering within four biological replicates, with a clear separation between genotypes in which HIF-1 is active versus inactive (Extended Data Fig. 3b). We compared the resulting transcriptional profiles (FDR<0.01; Supplemental Table 2) in four distinct pairwise combinations (active vs. inactive), deeming the intersection of those comparisons as high-stringency HIF-1-regulated genes (Extended Data Fig. 3a,c,d).

Direct targets of HIF-1 should (1) have a nearby HIF-1 binding site, and (2) be differentially expressed when HIF-1 is active. We adapted the BETA algorithm for *C. elegans* to integratively analyze our ChIP-seq and RNA-seq data^36^. Using a 15-kb window and an FDR<0.01, we identified 267 HIF-1 binding sites flanking 145 direct target genes (Fig. 1d; Extended Data Fig. 3e; Supplemental Table 3). Every identified direct target was exclusively upregulated rather than downregulated when HIF-1 was active, consistent with HIF-1 functioning as a transcriptional activator (Fig. 1e). Motif analysis of the direct targets using MEME-suite^37^ showed enrichment for the conserved HRE with a preference for “A” at position 15 (Fig. 1f). BETA identified *nhr-57* and *cysl-2* (Fig. 1g), two known HIF-1 targets, further validating our approach^*25,26,31,32*^. About half (53%) of the HIF-1 targets associated with a single binding site, and the magnitude of expression for targets loosely correlated with their number of binding sites (Fig. 1g). Interestingly, the genes *egl-9* and *rhy-1*, two key negative regulators of HIF-1^25^, contained some of the highest numbers of HIF-1 binding sites (4 and 8, respectively) and were dramatically upregulated by HIF-1 (10-fold and 76-fold, respectively), indicating a strong negative feedback loop and highlighting the importance of modulating the pathway itself as part of the response (Extended Data Fig. 3f).

### Characterizing HIF-1 Binding Sites

To determine if the HIF-1 binding sites for these targets are functional, we employed a transgenic approach. We examined one novel target, *pck-1* (PEP carboxykinase), associated with a single HIF-1 binding site, and one established target, *rhy-1*, associated with multiple binding sites. We generated fluorescent Venus transcriptional reporter transgenes containing genomic sequences encoding the HIF-1 binding site and the TSS (Fig. 2a; Extended Data Fig. 4a) fused to the sequence for Venus. In the case of *rhy-1*, our transgene only included the closest HIF-1 ChIP-seq peak upstream of the TSS. To quantify expression, we introduced into the genome a *Pmyo-2::mCherry* transgene in conjunction with the *pck-1::Venus* or *rhy-1::Venus* transgenes. As *myo-2* expression did not vary in our RNA-seq data sets, we normalized Venus expression to mCherry expression in the pharynx. Transgenic animals for both reporters had elevated levels of Venus fluorescence when HIF-1 was activated in *egl-9* mutants relative to wild type and *egl-9; hif-1* double mutants, where is HIF-1 is inactive (Fig. 2b-g; Extended Data Fig. 4b). We also generated versions of the transgenes in which either the complete sequence encoding the HIF-1 ChIP-seq peak was removed (ΔChIP), the 6-bp core HRE in the peak was deleted (ΔHRE), or only a minimal promoter sequence was present (Fig. 2a; Extended Data Fig. 4a). Removal of the full sequence encoding the ChIP-seq peak or the HRE alone was sufficient to reduce reporter expression in *egl-9* mutants to levels similar to that of the full-length promoter in wild type or in *egl-9; hif-1* double mutants (Fig. 2h; Extended Data Fig. 4b). These data show that HREs are required *in vivo* for HIF-1 to regulate target gene expression.

**Figure 2.**
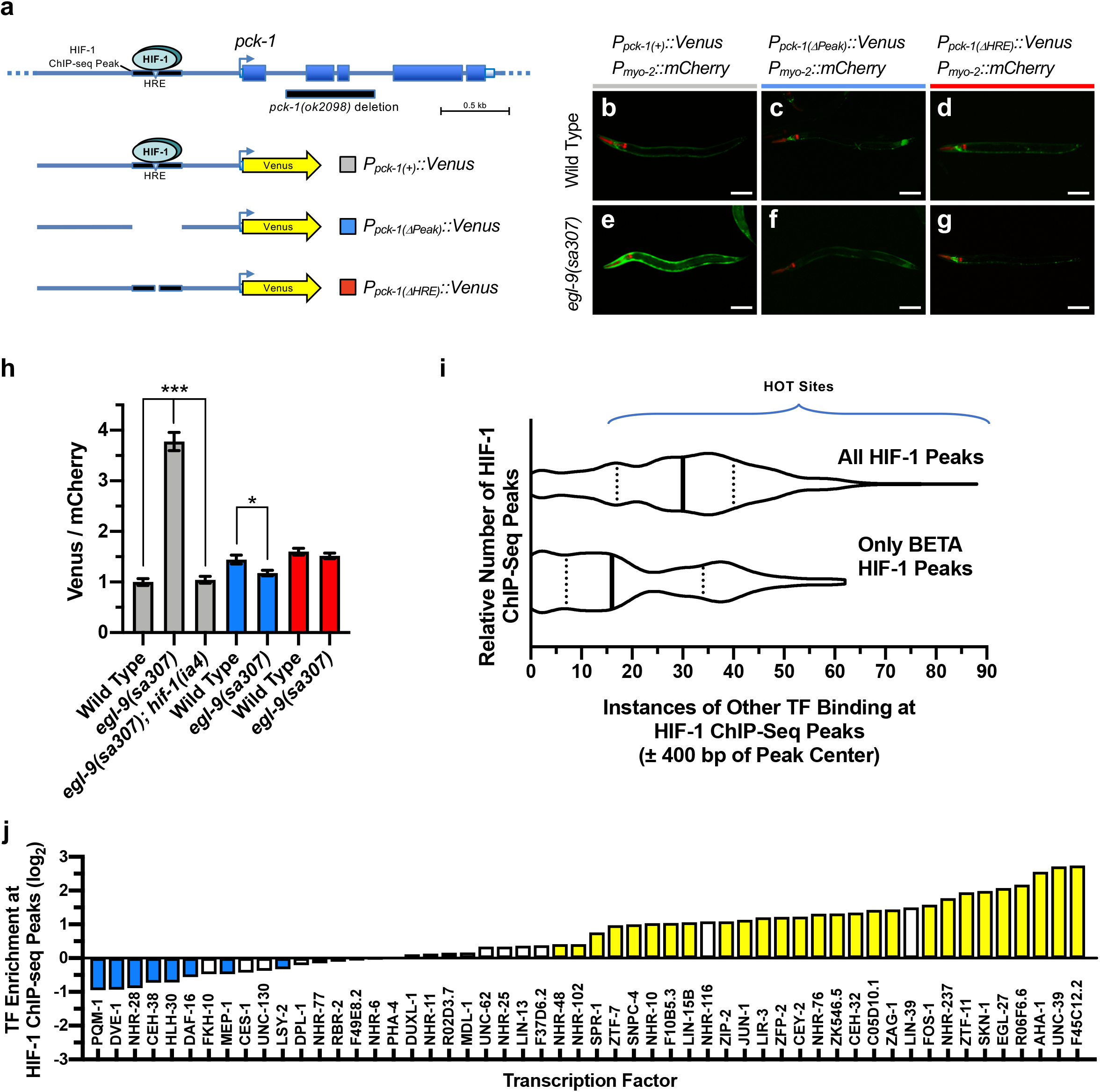
Characterizing HIF-1 Binding Sites. (a) Diagram of the *pck-1* locus and the different promoters used to drive Venus expression. The oval indicates HIF-1 and the black line beneath it indicates the HIF-1 binding site identified by ChIP-seq. The inverted triangle indicates the site of the HRE motif. Arrows indicate the TSS. Boxes indicate exons. The yellow arrow indicates sequences encoding the fluorescent Venus reporter. The black bar under the gene indicates sequences removed in the *pck-1(ok2098)* deletion. (b-g) Fluorescence images of animals expressing the indicated transgenes and with the indicated genotype. Venus fluorescence is indicated in green. Fluorescence from mCherry is red. Bar indicates 100 microns. (h) Graph of Venus/mCherry fluorescence ratios for the indicated genotype. Bar color indicates specific reporter as per panel A. Error bars indicate SEM. ***p<0.001, *p<0.05 ANOVA/Bonferroni Multiple Comparison test for the indicated comparisons. (i) Violin plot of histogram counting the number of HIF-1 binding sites (ChIP-seq peaks) against the number of other transcription factors (TFs) known to bind to each site’s region of the genome (within 400 bps). The top violin plot indicates all HIF-1 binding sites, whereas the bottom violin plot indicates only the functional HIF-1 binding sites identified by BETA. Solid line indicates median, whereas dotted lines indicate quartiles. HOT sites (peaks near the binding site of 15 or more other transcription factors) are indicated by the bracket. (j) Graph of the fold change in enrichment (log2) of each indicated transcription factor bound within 400 bps of a HIF-1 binding site (ChIP-seq peak center). TFs with overrepresented or underrepresented binding near HIF-1 sites compared to chance (p<0.01 via bootstrap confidence interval) are highlighted in yellow or cyan, respectively.

We reasoned that HIF-1 might share target genes with other stress response pathways such that other transcription factors might bind near or in HIF-1 binding sites. In addition, published ChIP-seq studies have identified High Occupancy Target (HOT) sites where occupancy by multiple (>15) transcription factors occurs; it is unclear whether HOT sites represent functional binding sites or simply “sticky” regions of the genome that lend to false signals within ChIP-seq data^38,39^. We surveyed the known binding sites of all transcription factors within the ModENCODE/ModERN database for L4 stage animals and determined which of the 55 factors also bound in or near HIF-1 sites. An examination of all 604 ChIP-seq peak sequences revealed the vast majority of them to be in or near known HOT sites (Extended Data Fig. 4c). By contrast, less than half of the 267 functional ChIP-seq peaks identified using RNA-seq data and BETA analysis were at HOT sites, suggesting many HOT sites might not be functional. Histograms of both sets of peaks (Fig. 2i) revealed three populations of HIF-1 binding sites: low occupancy (1-4 other factors bound), medium occupancy (5-20 other factors bound), and high occupancy (21 or more factors bound). Filtering by BETA caused a large drop in the number of high occupancy sites but only a subtle drop in low and medium occupancy sites, suggesting that the definition of HOT sites might need to be revisited. We observed enrichment for AHA-1 binding sites near HIF-1 sites (Fig. 2j; Extended Data Fig. 4d), as expected for a factor that acts as a HIF-1 dimerization partner^23^. We also observed enrichment for SKN-1 sites near HIF-1 sites, the *C. elegans* Nrf2 ortholog that promotes an antioxidant response^40^, suggesting shared targets between the hypoxia and antioxidant responses. By contrast, PQM-1 sites were underrepresented near HIF-1 sites, consistent with the increased hypoxia survival observed in *pqm-1* mutants^41^. We also note that, depending on the transcription factor, shared targets might be co-regulated, regulated synergistically, or regulated antagonistically.

### HIF-1 Reprograms Metabolism

To determine the types of genes regulated by HIF-1, we performed a Gene Ontology (GO) term analysis. For the 145 direct targets upregulated by HIF-1, we observed enrichment of GO terms for glycolysis, gluconeogenesis, amino acid metabolism, sulfur oxidation, fatty acid beta-oxidation, and oxidation-reduction metabolism (Fig. 3a). The remaining targets that were indirectly upregulated by HIF-1 were enriched for deoxyribonucleotide metabolism and innate immunity (Extended Data Fig 5a). The increased expression of innate immunity genes could explain changes in host-pathogen interaction observed in hypoxia response pathway mutants^42,43^. The targets indirectly downregulated during HIF-1 activation were enriched for GO terms for protein dephosphorylation, multicellular development, transcriptional regulation, organelle organization and biogenesis, signal transduction, and nucleic acid metabolism (Extended Data Fig 5b). Taken together, these results suggest that HIF-1 reprograms metabolism by directly binding to the promoters of key metabolic pathway enzymes, promoting their expression.

**Figure 3.**
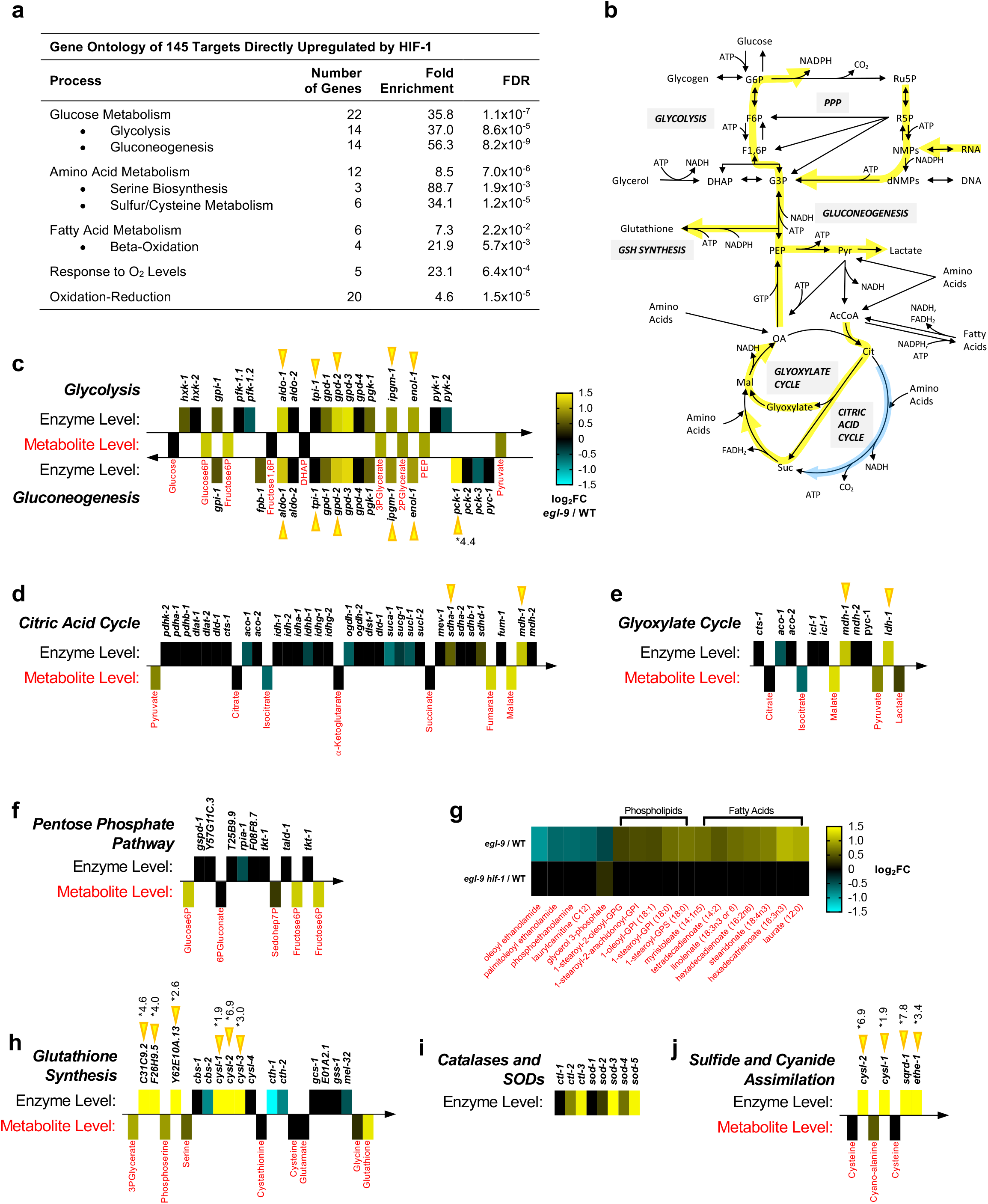
HIF-1 Reprograms Metabolism. (a) Enriched Gene Ontology (GO) classifications for the 145 direct HIF-1 targets. Fold enrichment and FDR for each classification and subclassification is indicated. (b) Overview of major metabolic pathways impacted by HIF-1 activation. Yellow and cyan arrows indicate pathways upregulated or downregulated, respectively, in *egl-9* mutants relative to wild type. (c-j) Combined heat map for metabolites (red) and the enzymes (black) and directional pathways (arrows) known to regulate their metabolism. On one side of each pathway arrow, differentially colored boxes indicate enzyme mRNA expression levels (log_2_ fold change) in *egl-9* mutants relative to wild type; enzymes are labeled in black by their indicated *C. elegans* gene name. On the other side of each pathway arrow, differentially colored boxes indicate metabolites levels (log_2_ fold change) in *egl-9* mutants relative to wild type (and *egl-9; hif-1* double mutants relative to wild type in panel g); metabolites are labeled in red. Gene expression changes that exceed a log_2_ fold change of 1.5 are marked with an asterisk showing the actual log_2_ fold change value. Yellow arrow heads indicate genes with HIF-1 binding sites and showing direct regulation by HIF-1. Glycolysis and gluconeogenesis are graphed together, in opposite direction, as they share multiple metabolites and enzymes.

To confirm that activated HIF-1 reprograms metabolism, we used a metabolomics approach to determine if the transcriptionally upregulated pathways were more populated with metabolites from those pathways. We used liquid chromatography and mass spectrometry (LC-MS) to obtain metabolomic profiles of L4 stage wild type and *egl-9(sa307)* mutants under aerobic conditions, examining 558 metabolites. We observed differences (p<0.05) in the levels for 175 metabolites (Supplemental Table 4). HIF-1 activation led to elevated levels of various amino acids, carbohydrates, lipids, and nucleotides, suggesting increased flux through multiple metabolic pathways (Fig. 3b).

To examine metabolite levels in the context of specific biochemical pathways, we mapped wild type versus *egl-9* mutant metabolome data against corresponding RNA-seq data for enzymes that catalyze reactions involving each specific metabolite. It is well established that HIF-1 activation in mammals promotes glycolysis by upregulating several key enzymes in the pathway^43,44^. Consistent with these mammalian studies, we found that HIF-1 directly promoted the expression of multiple key glycolysis enzymes and increased metabolite population of this pathway (Fig. 3c), highlighting a conserved function of HIF-1 activation between nematodes and mammals. Glycolysis generates pyruvate, which enters the tricarboxylic acid (TCA) cycle to generate NADH under aerobic conditions. Oxygen is required to oxidize that NADH and thereby generate ATP; thus, we would expect metabolite flux in the TCA cycle to be reduced when HIF-1 is activated.

As expected, most TCA cycle enzymes and metabolites are unchanged in *egl-9* mutants (Fig. 3d). However, *egl-9* mutants had elevated levels of *mdh-1* (malate dehydrogenase) and *ldh-1* (lactate dehydrogenase), as well as associated metabolites malate, pyruvate, and lactate (Fig. 3e). Elevated malate and malate dehydrogenase are indicative of flux through the glyoxylate cycle, a variation of the TCA cycle used in some organisms to convert citrate to oxaloacetate to feed the gluconeogenesis pathway.

We also observed indicators of increased gluconeogenesis (Fig. 3c). As previously discussed, HIF-1 directly and dramatically promotes the expression of *pck-1* (PEP carboxykinase), which converts oxaloacetate (OA) to phosphoenolpyruvate (PEP), a key rate-limiting step specific to gluconeogenesis. In addition to regenerating glucose, gluconeogenesis provides substrates for the pentose phosphate pathway (PPP), which generates reducing equivalent NADPH, 5-carbon sugars (e.g., ribose-5-phosphate used to synthesize nucleic acids), and erythose-4-phosphate (used to synthesize aromatic amino acids). We observed an mincrease in PPP metabolites in *egl-9* mutants relative to wild type (Fig. 3f), although none of the key enzymes within the PPP were direct targets of HIF-1, suggesting that increased PPP flux is an indirect effect of HIF-1 activation of gluconeogenesis.

With elevated PPP flux, we would expect that downstream pathways that utilize NADPH to be active, with enrichment of associated metabolites in *egl-9* mutants compared to wild type. For example, NADPH generated from PPP serves as reducing equivalent for fatty acid synthesis, and both synthesized and dietary fatty acids are attached to glycerol-3-phosphate (G3P) for conversion to storage lipids. Consistent with metabolic pathway, we observed decreased G3P and increased levels of certain fatty acids when HIF-1 is active (Fig. 3g). Phospholipids are also synthesized from G3P, and we found that active HIF-1 increased the levels of certain phospholipid species, as well as endocannabinoid-like fatty acid ethanolamides and lysophospholipids often used as secondary signaling molecules (Fig. 3g).

NADPH also replenishes reduced glutathione, a major antioxidant for combating oxidative stress, and we observed higher levels of glutathione in *egl-9* mutants relative to wild type (Fig. 3h). Indeed, many of the direct target genes most dramatically upregulated by HIF-1 lie within pathways that produces glutathione (Fig. 3h), and the catalases and superoxide dismutases that combat oxidative stress are all indirectly upregulated (Fig. 3i). Oxidative stress response pathways help fight bacterial infection in *C. elegans*, as bacterial pathogens like *Pseudomonas aeruginosa* produce toxins like HCN, which mimics hypoxia, and H2S^44,45^. We observed that HIF-1 directly promotes the expression of the entire H2S/HCN detoxification pathway (Fig. 3j), including *cysl-2* (cyanoalanine synthase) and *sqrd-1* (sulfide quinone oxidoreductase), which convert these toxins to polysulfides and sulfates^44^. Upregulation of H2S/HCN detoxification by HIF-1 is consistent with *egl-9* and *hif-1* mutants being resistant and sensitive, respectively, to *Pseudomonas* infection and sulfide/cyanide toxicity.

### HIF-1 and PCK-1 Are Required for Adaptive Survival to Hypoxia and Oxidative Stress

Traditionally, HIF-1 is thought to protect against hypoxic stress by promoting anaerobic ATP synthesis, yet our multiomics analyses highlighted its promotion of the oxidative stress response. Gluconeogenesis plays a central role in promoting oxidative stress resistance through generation of NADPH and glutathione. We therefore examined null mutants for *pck-1*, the rate limiting enzyme of gluconeogenesis directly regulated by HIF-1. These *pck-1(ok2098)* mutants were viable under aerobic conditions, most likely because two other PEP carboxykinases, *pck-2* and *pck-3*, compensate for baseline function. Neither *pck-2* nor *pck-3* showed HIF-1-dependent regulation, suggesting *pck-1* is a specific HIF-1-induced isoform. We examined changes in oxidative stress sensitivity directly by exposing L4-stage animals to a lethal dose of paraquat, a superoxide generator. Less than half of wild-type animals survive after 15 hours of paraquat exposure. By contrast, most *egl-9* mutants survived, and this survival was HIF-1- and PCK-1-dependent (Fig. 4a). PCK-1 mediates gluconeogenesis by producing PEP, and we observed elevated PEP levels when HIF-1 was activated in *egl-9* mutants (Fig. 3c). Supplementation of animals with PEP was sufficient to restore survival to *hif-1* and *pck-1* mutants exposed to paraquat (Fig. 4a), demonstrating the importance of PCK-1 and gluconeogenesis upregulation by HIF-1 in combating oxidative stress.

**Figure 4.**
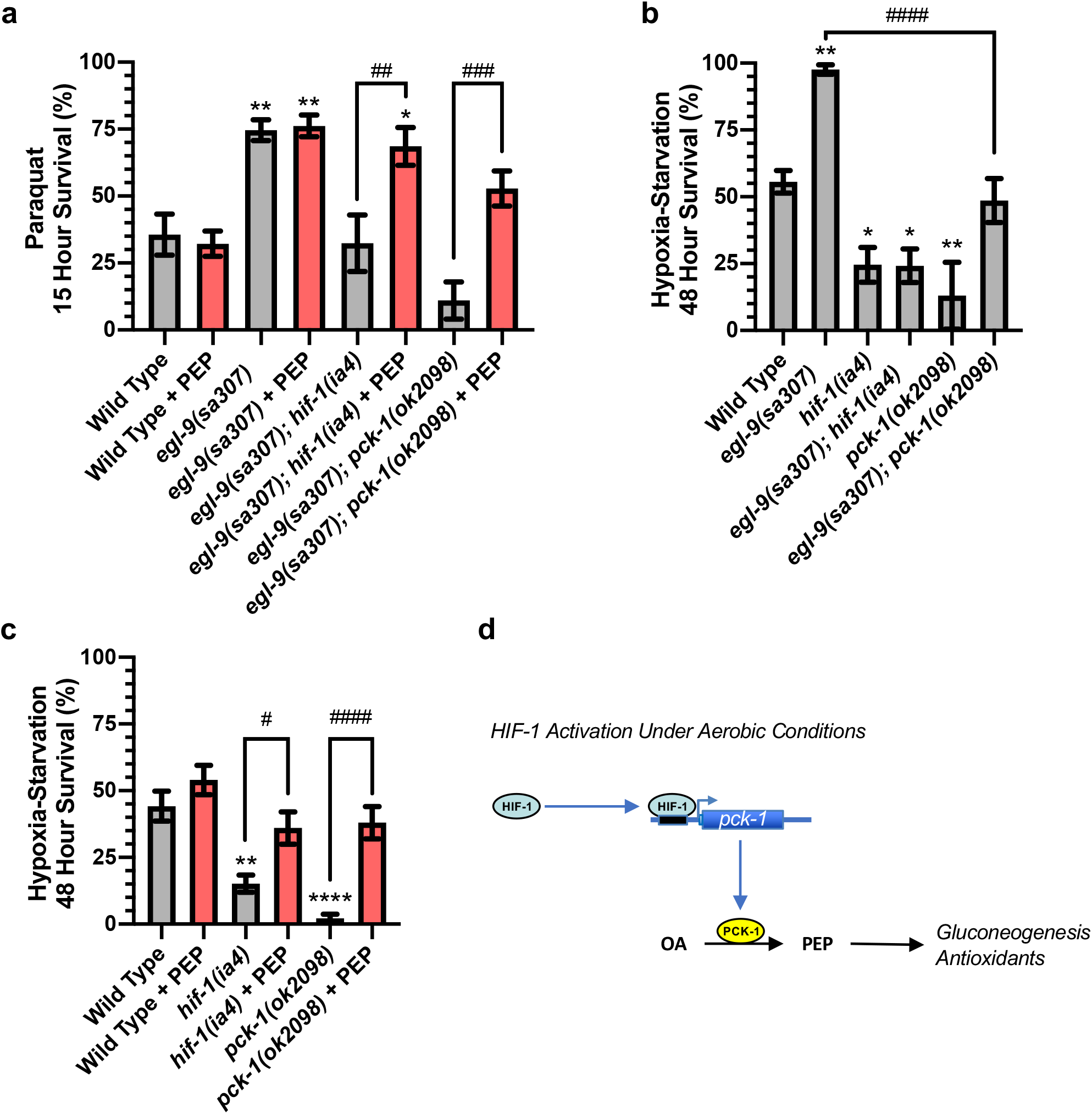
HIF-1 Targets Required for Adaptive Survival. (a) Percent of L4-stage animals to survive paraquat (200 mM, 15 hours, 20°C, on plates with food). (b,c) Percent of L4-stage animals to survive hypoxia-starvation (0.5% O_2_, 48 hours, 20°C, in liquid without food). Animals previously grown on plates supplemented with 50 mM PEP are indicated by red bars. ****p<0.0001, **p<0.01, *p<0.05 ANOVA/Dunnett’s multiple comparison test to wild type. ####p<0.0001, ###p<0.001, ##p<0.01, #p<0.05 ANOVA/Sidek’s multiple comparison test for the indicated comparisons. (d) Model illustrating how HIF-1 binds near the *pck-1* promoter to upregulate PCK-1 expression and drive gluconeogenesis and the production of antioxidants.

Given that both hypoxia and subsequent reoxygenation result in the production of reactive oxygen species (ROS) and oxidative stress, we reasoned that HIF-1 and PCK-1 might also protect against hypoxia. We exposed L4-stage animals to 48 hrs of hypoxia without food and measured survival after 24 hrs of recovery at normoxia. While only half of wild-type animals survived this assay, nearly all *egl-9* mutants survived, and this survival was HIF-1- and PCK-1-dependent (Fig. 4b). In addition, both *hif-1* and *pck-1* single mutants were significantly susceptible to hypoxia, and this susceptibility was rescued by PEP supplementation (Fig. 4b,c). Although more variable by trial, we observed similar results with a hypoxia survival assay on food at higher temperature (Extended Data Fig. 6a,b). We also observed that *gpd-3(ok2870)* mutants, which are incapable of continuing gluconeogenesis, survived hypoxia poorly (Extended Data Fig. 6a,b). Mutations in *pck-1* did not suppress the egg retention or extended lifespan phenotypes of *egl-9* mutants (Extended Data Fig. 6c,d), indicating that the upregulation of PCK-1 does not mediate all functions of HIF-1. Taken together, our results suggest that HIF-1 activation allows animals to survive hypoxia by primarily and directly promoting gluconeogenesis and oxidative stress resistance.

## Discussion

Transcriptional effectors of stress response pathways coordinate the regulated expression of multiple target genes needed to rebalance physiological homeostasis over time and in different tissues. As an effector for both the hypoxia response pathway and the Warburg effect, HIF-1 promotes ATP synthesis by promoting glycolysis, yet the nature and role of its full array of targets *in vivo* is not understood. Our studies combined ChIP-seq and RNA-seq to identify a full array of direct HIF-1 targets, combined with metabolomics to understand the physiological changes resulting from their expression. We also observed changes in expression of indirect targets, which could reflect either a second phase of regulation by transcription factors elevated during the first phase or the homeostatic action of other physiological pathways responding to changes due to HIF-1 activation. Our metabolomics analysis confirmed that the HIF-1-mediated expression of these targets reprograms metabolism beyond simply augmenting glycolysis Instead, we found that HIF-1 directly promotes lipid metabolism, the glyoxylate cycle, gluconeogenesis, the PPP, H2S/HCN detoxification, and glutathione synthesis, making ours the first study to combine genomic, transcriptomic, and metabolomic approaches to characterize the action of HIF-1 activation *in vivo*

Activation of the glyoxylate cycle was intriguing, as it allows some organisms to mobilize lipids into acetyl-CoA, which can then be converted into oxaloacetate for gluconeogenesis and/or lactic acid production. Compared to the TCA cycle, the glyoxylate cycle does not produce as much NADH requiring oxidation nor carbon lost as CO2, which would be advantageous under hypoxic conditions. *C. elegans* could use this cycle to derive energy from stored lipids under fairly anaerobic conditions when beta oxidation would not be effective. It could also be used to derive glucose from stored lipids, which could be used to generate reducing power, antioxidants, or macromolecules via the PPP. Although the glyoxylate cycle was assumed to be absent in vertebrates, more recent evidence suggests that a vertebrate version of the pathway acts in the liver^46,47^.

HIF-1 directly promotes the expression of the *pck-1* PEP carboxykinase, a key mediator of gluconeogenesis, as well as metabolic flux through the pathway (Fig. 4d). Gluconeogenesis provides the carbon needed to make glutathione and produce NADPH reducing equivalents required to combat ROS and oxidative stress. Indeed, we find that *pck-1* mutants survive hypoxic stress as poorly as *hif-1* mutants, and HIF-1 activation can offset damage by agents that cause oxidative stress. Our results highlight that the need for HIF-1 activation in mobilizing antioxidant defenses is likely as important as its role in promoting anaerobic energy production. Moreover, blocking gluconeogenesis might present an approach to reduce progression of HIF-1-positive tumors.

## METHODS

### Generation of Transgenes and Transgenic Animals

The GFP-tagged *hif-1* transgene was obtained as a genomic fosmid from the TransgeneOme Project^30^. The polyclonal stab culture was streaked out under triple selection (chloramphenicol, streptomycin, nourseothricin) and individual clones were selected for transgene construct validation by sequencing. The construct was then stably integrated into the genome of *unc-119(ed3)* animals using microparticle bombardment^48^. The resulting stably integrated line *odIs131* was made homozygous by selecting single hermaphrodites that gave 100% Non-Unc progeny, then outcrossed four times. Subsequently, the *unc-119(ed3)*; *odIs131* strain was crossed to *unc-119(ed3); egl-9(sa307); hif-1(ia4)* to generate the following strains used for ChIP-seq.

To generate *Ppck-1::Venus* transgenic animals, 2816 bp upstream of the ATG of *pck-1* was amplified and cloned upstream of a *venus* coding sequence present in pPD95.77-mVenus. Using Q5-site directed mutagenesis (Life Technologies Ltd), the ΔPeak (1084 – 912 upstream of start) and the ΔHRE (1019 – 1003 upstream of start) were deleted to yield *Ppck-1(ΔPeak)::Venus* and *Ppck-1(ΔHRE):Venus*, respectively. These three plasmids (100 ng/μL) were injected into wild-type animals at 100 ng/μL along with a *Pmyo-2::mCherry* (50 ng/μL) co-injection marker. At least two independent lines were tested for each experiment.

To generate *Prhy-1::Venus* transgenic animals,1510 bp upstream of the ATG of *rhy-1* was amplified and cloned upstream of a *venus* coding sequence as above. Q5-site directed mutagenesis (Life Technologies Ltd) was used for the removal of the ΔPeak (1170 – 617 bp upstream of start), ΔHRE1 (971 – 956 bp upstream of start), ΔHRE2 (946 – 931 bp upstream of start) and *min* (591 p to start) variants of the promoter. Each plasmid (100 ng/μL) was injected into wild-type animals at 100 ng/μL along with a *Pmyo-2::mCherry* (50 ng/μL) co-injection marker. At least two independent lines were tested for each experiment.

### Epifluorescence Microscopy and Image Analysis

Fluorescent proteins were visualized in nematodes by mounting on 2% agarose pads with 10 mM tetramisole. All animals were synchronized by alkaline bleaching and visualized at the L4 stage. Fluorescent images of transgenic animals containing the *odIs131[hif-1::gfp], Ppck-1::Venus, Prhy-1::Venus*, and/or *Pmyo-2::mCherry* transgenes were observed using an AxioImager M1m (Carl Zeiss, Thornwood, NY). A 10X or 40X PlanApo objective was used to detect fluorescence. Images were acquired with an ORCA charge**‐**coupled device camera (Hamamatsu, Bridgewater, NJ) by using iVision software (Biovision Technologies, Uwchlan, PA). Exposure times were chosen to capture at least 95% of the dynamic range of fluorescent intensity of all samples. Images were quantified in Fiji by selecting each entire animal using the segmented line tool and measuring pixel intensity for the different channels^49^. All data with normal distributions were analyzed with GraphPad Prism 8 in most cases using ANOVA with Dunnett’s post-hoc test correction for multiple comparisons.

### Quantitative RT-PCR to Assess HIF-1::GFP mRNA Expression

Two sets of primers were used to assess the expression level of the endogenous *hif-1* as well as the *odIs131[hif-1::gfp]* transgene. The first set (5’-ACTTGCCTGACTTTACACCTG-3’ and 5’-TGTTGGAATGGTTGATAATGTTGAG-3’) amplified a sequence in the 3’ end of all *hif-1* mRNA transcripts. The second set (5’-GCTTGGACGGCTTTGTTATG-3’ and 5’-GAAGGGCTCGACCTGTTAAAT-3’) amplified a sequence within the *hif-1(ia4)* deletion. Quantitative RT-PCR was performed on mRNA isolated from L4 stage animals for the relevant genotypes, and relative mRNA abundance was performed using *act-5* as a normalization control.

### Characterizing Egg Laying and Egg Retention

All genotypes were synchronized by alkaline bleaching and arrested at L1 stage overnight in M9 buffer. Synchronized genotypes were assayed 43-46 hours after reaching the L4 stage, when embryos present within adult animals were counted using a dissection microscope.

### ChIP-seq

ChIP-Seq was performed on L4 stage by the modERN/modEncode consortium as per their standard protocol^38^. Developmental synchronization was achieved by bleaching and L1 arrest. Arrested L1s were plated on NGM plates seeded with OP50 bacteria and grown for 6 hr at 20°C for L1 collection. L4 stage animals were harvested, and samples were cross-linked with 2% formaldehyde for 30 min at room temperature and then quenched with 1 M Tris pH 7.5. The pelleted nematodes were subsequently flash frozen in liquid nitrogen and stored at -80°C. Pellets were thawed on ice and 750 ml of FA buffer containing protease inhibitors (Roche Cat#11697498001 Complete Protease Inhibitor Cocktail Tablet, 125 µl 100 mM PMSF, and 25 µl 1 M DTT per 25 ml FA buffer) was added, and samples were then transferred to a 2 ml KONTES dounce (Kimble Chase, Vineland, NJ). Samples were dounced on ice 15 times with the small “A” pestle for two cycles with a 1 min hold between each cycle. Samples were then dounced 15 times with the large “B” pestle for four rounds with a 1 min hold between each cycle. Samples were then sonicated to shear chromatin into 200–800 bp DNA fragments.

For each sample, 4 mg of protein lysate was immunoprecipitated using anti-GFP antibodies (gifts of Tony Hyman and Kevin White). GammaBind G Sepharose beads (GE Healthcare Life Sciences) were pre-washed and blocked in binding buffer and 0.1 mg/ml BSA. Samples were pre-cleared by adding 100 µl of 50/50 bead/buffer solution at 4°C, followed by centrifugation. Samples of each replicate were removed and pooled to serve as total chromatin input. To each replicate, 15 µg of antibody was added overnight at 4°C, followed by another overnight of incubation with the bead mix. Immunoprecipitates (IPs) were washed four times with cold lysis buffer and twice with cold TE. Pellets were resuspended in elution buffer (10 mM EDTA, 1% SDS, and 50 mM Tris-HCl pH 8) and incubated at 65°C for 10 min. Samples were centrifuged and supernatants transferred to a fresh tube. Pellets were resuspended in 29% TE and 0.67% SDS and immediately centrifuged. Elution supernatants were combined and incubated at 65°C with mild shaking overnight. Chromatin input samples were incubated at 60°C with mild shaking overnight following the addition of Proteinase K and SDS to final concentrations of 0.1 mg/ml and 0.01%, respectively. The next day, inputs were incubated at 70°C for 20 min. Proteinase K was added to each IP and incubated at 50°C for 2 hr. RNaseA was added to the chromatin input to a concentration of 0.017 mg/ml and incubated at 37°C for 2 hr. DNA was purified with MinElute columns (QIAGEN, Valencia, CA), eluting in 13 µl (elution buffer provided with MinElute kit). An additional 48 µl EB was added to input samples after purification. Samples were stored at -20°C.

The enriched DNA fragments and input control (genomic DNA from the same sample) for two biological replicates were used for library preparation and sequencing. Samples were libraried and multiplexed using the Ovation Ultralow DR Multiplex Systems 1–8 and 9–16 (NuGEN Technologies, San Carlos, CA) following the manufacturer’s protocol, except that QIAGEN MinElute PCR purification kits were used to isolate the DNA. Briefly, 1 µl of input DNA and 10 µl of IP DNA was used to prepare sequencing libraries using NuGEN Ultralow library kits. Samples were prepared according to the manufacturer’s protocol. Sequencing was performed on the Illumina HiSequation 2000/2500/4000.

The Illumina sequencing data were aligned to the reference genome using the Burrows– Wheeler Aligner (BWA). Data were aligned to genome version WS245. Peak regions significantly enriched in aligned reads were called by ChIP-seq processing pipeline standard for modERN/ModENCODE^38^. Peaks above an irreproducibility discovery rate (IDR) of 0.1% were used to generate final peak sets.

ChIP-seq data sets are available at NIH/NCBI GEO through accession number GSE7173333.

### RNA-seq

Developmentally synchronized animals were obtained by hypochlorite treatment of gravid adults and embryos hatched overnight for 15-17 hours in M9. Starvation-arrested L1s were plated on NGM plates and grown at 20°C until L4 stage. Total RNA was isolated from animals using Trizol (Invitrogen) combined with Bead Beater lysis in four biological replicates for each genotype. An mRNA library (single-end, 50-bp reads) was prepared for each sample/replicate using Illumina Truseq with PolyA selection (Genewiz or RUCDR). Libraries were sequenced across two lanes on an Illumina HiSeq2000 (GeneWiz) or an Illumina HiSeq 2500 in Rapid Run Mode (RUCDR). Reads were mapped to the *C. elegans* genome (WS245) and gene counts generated with STAR 2.5.1a. Normalization and statistical analysis on gene counts were performed with EdgeR using generalized linear model functionality and tagwise dispersion estimates. Likelihood ratio tests were conducted in a pairwise fashion between genotypes with a Benjamini and Hochberg correction. Genes were considered to be HIF-1-dependent if they were differentially expressed with an FDR<0.01 in the same direction (up or down) in all four of the following pairwise comparisons: (1) *egl-9* vs. N2; (2) *egl-9* vs. *egl-9 hif-1*; (3) *odIs131; egl-9* vs. N2; (4) *odIs131; egl-9* vs. *egl-9 hif-1*. RNA-seq data sets are available at NIH/NCBI GEO through accession number GSE173581.

### Identification of direct targets using BETA basic

BETA basic was used to identify potential direct targets for HIF-1 for the WS245 annotation of the *C. elegans* genome^36^. The following parameters were used: 15kb from TSS, FDR cutoff of 0.01 and one-tail KS test cutoff of 0.01. The input files consisted of .bed files of IDR thresholded peaks and differential expression Log2FC and FDR values for the different genotypic pairwise comparisons. The final list of targets was the intersection of these comparisons.

### Motif Identification and Enrichment

The sequences for the middle of each ChIP-Seq peak (+/-100 bp, repeat masked with N) for were extracted from the UCSC Genome Browser and entered into the MEME-Chip tool at meme-suite.org^37^. The following parameters were used: JASPA2018 Core Vertebrate Database (non-redundant), motif width from 6-15 bp, 1^st^ order background model, STREME cutoff of 0.05.

### Identification of Transcription Factor Occupancy at HIF-1 Binding Sites

Binding site coordinates (IDR Thresholded peaks, FDR < 0.01) for all transcription factors for which ChIP-seq was conducted at the L4 stage were obtained from the modEncode repository^38^. In order to determine the number of instances these peaks overlapped with the HIF-1 ChIP-seq peaks for OR3350, we developed an R script that is available at https://github.com/shahlab/hypoxia-multiomics.

### Metabolomics

All genotypes were age-synchronized by alkaline bleaching, and arrested L1 larvae were plated on 100 mm plates containing standard NGM media with OP50. Animals were grown at 20°C until L4 stage, at which point they were washed into 50 ml conical bottom tubes and allowed to settle for 15 mins. Animals were then washed with 3 x 40 ml sterile M9 and centrifuged at 2000 rpm for 5 mins. Five hundred µL of the final pellet was transferred to a 15 ml conical bottom tube and flash frozen in liquid N2 and stored at -80 °C until sent to Metabolon (Metabolon Ltd) for metabolomic processing. Samples were extracted with methanol, and each extract was divided into four fractions: two for analysis by two separate reverse phase (RP)/UPLC-MS/MS methods with positive ion mode electrospray ionization (ESI), one for analysis by RP/UPLC-MS/MS with negative ion mode ESI, and one for analysis by HILIC/UPLC-MS/MS with negative ion mode ESI. Raw data was extracted and peak-identified, then compared to a library based on authenticated standards.

### Paraquat Survival Assay

Paraquat sensitivity was performed according to the acute paraquat sensitivity assay described in Senchuk et al ^50^. Animals were placed at the L4 stage at 20°C and survival was counted every hour until all animals were dead. At least 20 animals per strain per trial were used for the assays.

### Hypoxia Survival Assays

For hypoxia-starvation, M9 solution was pre-equilibrated with 0.5% O2 for at least an hour. Nematodes were grown on standard NGM plates. Age-synchronized L4 animals (n ≥ 20) were then collected from plates, washed with M9 buffer, and then incubated with hypoxic M9 solution in capped tubes at 20°C for the indicated time. After hypoxia-starvation, nematodes were collected by centrifugate and returned to standard NGM plates under normoxia for 24 hours to allow recovery. Survival of animals was counted by assessing movement and response to gentle touch by a platinum wire pick.

For hypoxia on plates with food, a nitrogen-displacement hypoxia chamber (BioSpherix) was pre-equilibrated to 0.5% O2 for at least 2 hrs. Standard NGM plates were ringed with garlic extract then allowed to dry 30 minutes to prevent animals from escaping up the plate sides during hypoxic exposure^51^. Age-synchronized L4 animals (n ≥ 20) were then placed in the plate and incubated at 25°C, 0.5 % O2, for 48 hrs. Control plates were placed in 21.1 % O2 at 25°C. At the end of 48 hrs, plates were moved to 20°C for 24 hrs, after which survival of animals was counted by assessing movement and response to gentle touch by a platinum wire pick.

### PEP Supplementation

For the paraquat survival and hypoxia assays, nematodes were grown on standard NGM plates containing 50 mM PEP prior to collection at the L4 stage for the assay. The same protocol was then followed as above for each assay. No PEP was included in the incubation plates, M9 solution, or recovery plates.

## ACKNOWLEDGEMENTS

We thank the TransgenOme Project for the *hif-1::gfp* fosmid, Peter Schweinsberg and Barth Grant for help with bomdbardment integration, Daja O’Bryant and Hazel Schubert for assistance in establishing bioinformatic pipelines, and Michelle Kudron and Valerie Reinke for their assistance with the ChIP-seq. This work was supported by an NIH grant (R01GM101972) to CR and a New Jersey Commission on Spinal Cord Research Postdoctoral Fellowship (CSCR13FEL001) to SMP.

## AUTHOR CONTRIBUTIONS

MV, SMP, and CR designed the overall set of experiments, as well as analyzed and interpreted the data. SMP and TLM mediated the ChIP-seq and RNA-seq, whereas MV, SMP, PS, and CR analyzed the results. MV mediated and analyzed the metabolomics. MV, AP, and NSK constructed and analyzed the *pck-1* promoter mutations. MV and CR conducted hypoxia and oxidative stress survival assays, as well as wrote the manuscript.

## COMPETING INTEREST DECLARATION

The authors declare no competing interests.

## EXTENDED FIGURE LEGENDS

**Extended Data Figure 1.**
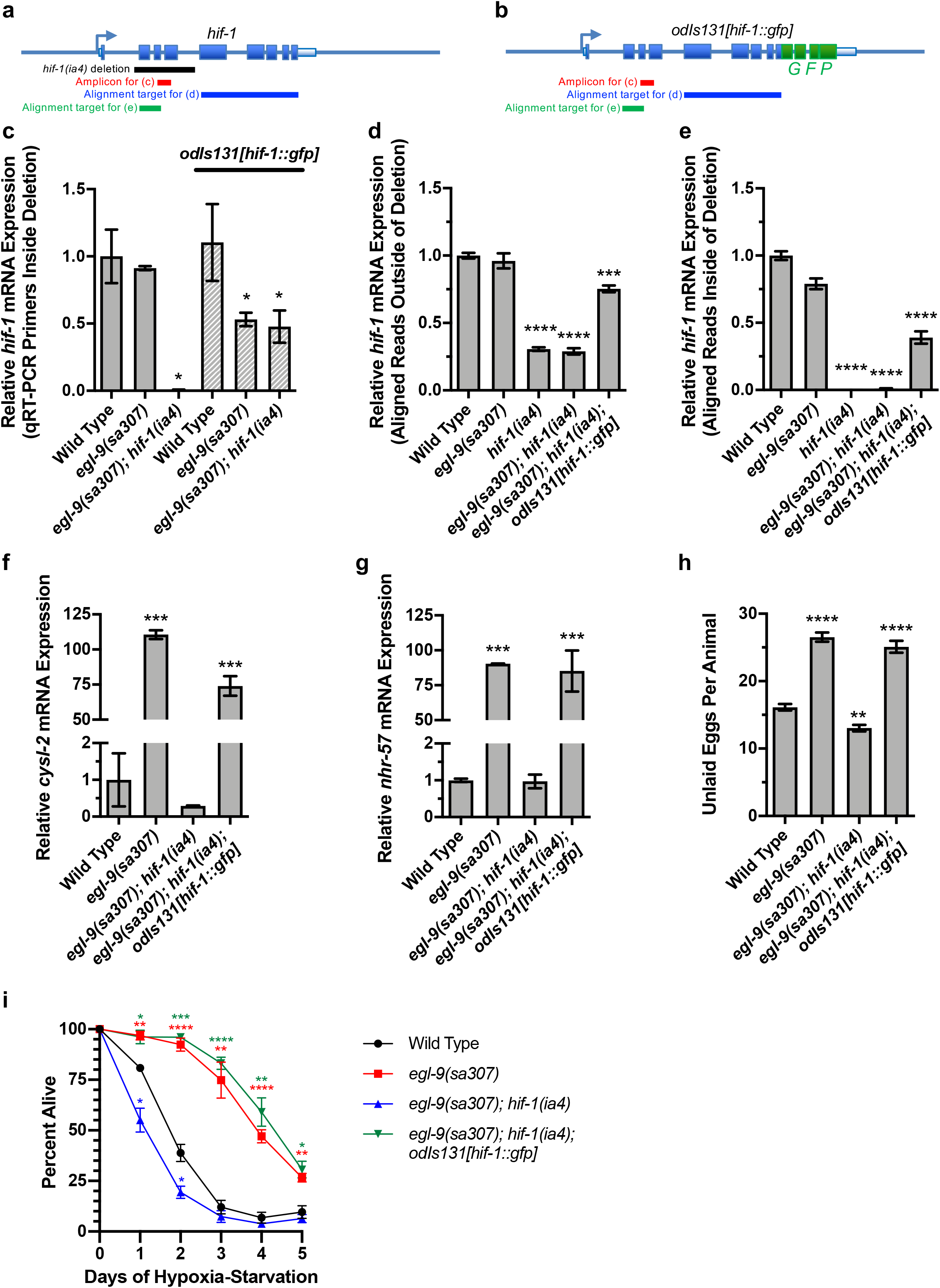
The *odIs131[hif-1::gfp]* Transgene Behaves Like Endogenous *hif-1*. Diagrams of the (a) *hif-1* locus and the (b) *odIs131[hif-1::gfp]* transgene. Arrows indicate the TSS. Boxes indicate exons. The black bar indicates sequences removed in the *hif-1(ia4)* deletion. The red bar indicates the amplicon for the qRT-PCR in (c). The blue and green bars indicate the sequence targets to which RNA-seq reads were aligned in (d) and (e), respectively. (c-e) Relative *hif-1* mRNA expression levels determined by (c) qRT-PCR or (d,e) RNA-seq read counts per total million reads in the indicated genetic background. Primers in (c) anneal to sequences inside the *hif-1(ia4)* deletion and thus only detect transgenic *hif-1* transcripts in the *egl-9(sa307); hif-1(ia4); odIs131[hif-1::gfp]* background. RNA-seq sequencing reads were aligned to (d) exons 5-9 outside of the deletion (thus measuring both wild-type transcripts and transcripts containing the deletion), or (e) exons 2-3 inside of the deletion (thus measuring only wild-type transcripts). The homozygous *odIs131[hif-1::gfp]* transgene inserted on the X chromosome; its expression level being about half that of endogenous *hif-1* is consistent with the transgene undergoing X-linked dosage compensation. All expression data is normalized to wild type in each graph. (f,g) Relative mRNA expression levels for the indicated HIF-1 target genes in the indicated genotypes determined by qRT-PCR. (h) Average number of unlaid eggs *in utero* per animal for the indicated genotype (n=40-55 animals). (i) Average percent of animals alive on the given day of exposure to hypoxia without food (n=5 trials, 50-120 animals per trial). Results in (d-g) demonstrate that the *odIs131[hif-1::gfp]* transgene is sufficient to conduct HIF-1 physiological functions when expressed at the level of endogenous *hif-1*, which is expected given that all reported *hif-1* mutant phenotypes are recessive. For all graphs, error bars indicate SEM. ****p<0.0001, ***p<0.001, **p<0.01, *P<0.05 compared to wild type using ANOVA/Dunnett’s multiple comparison test.

**Extended Data Figure 2.**
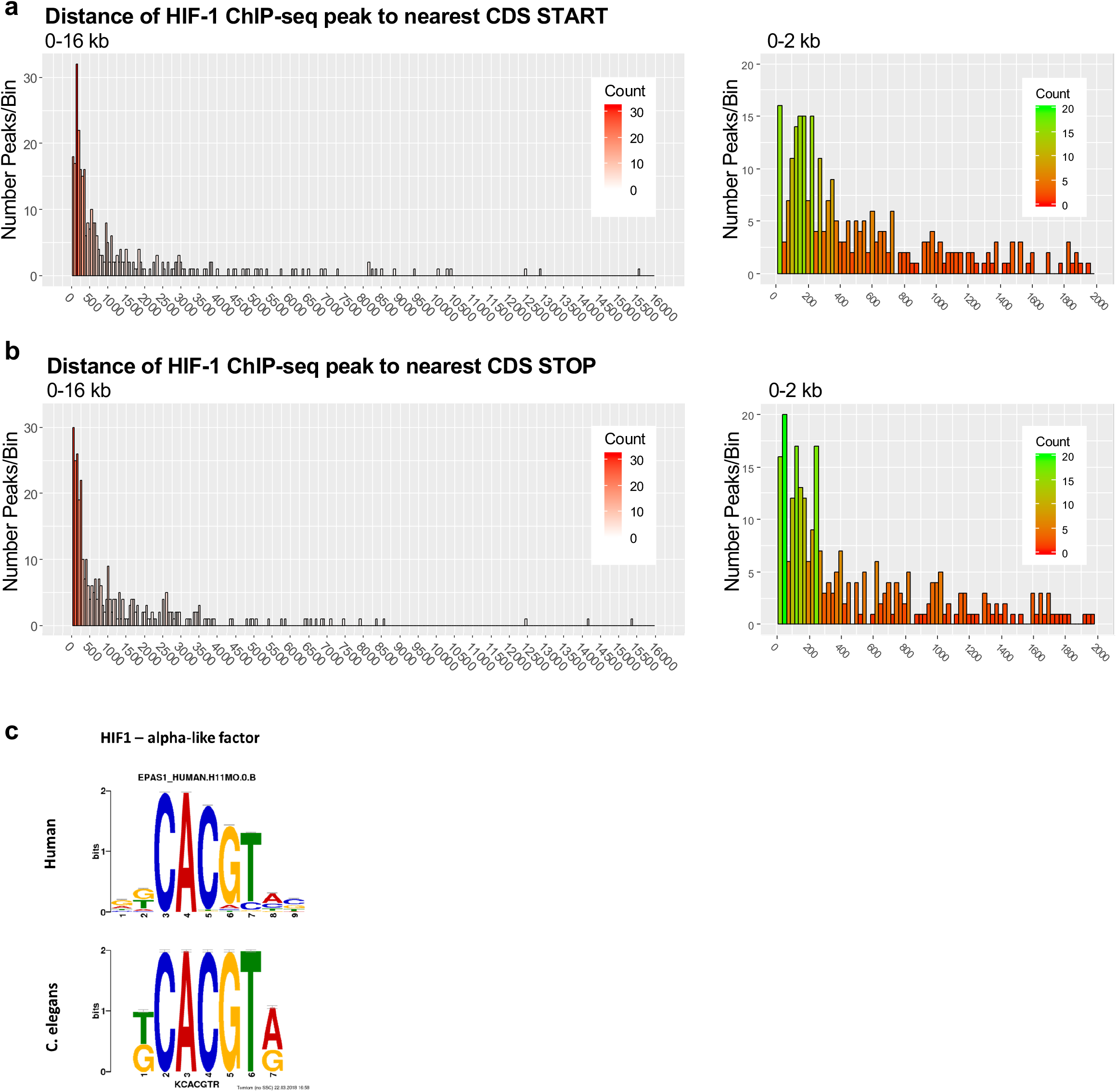
(a,b) Histogram of the number of HIF-1 binding sites (ChIP-seq peaks) against the distance of those peaks from (a) the TSS of the nearest gene, or (b) the stop codon of the nearest gene. Separate graphs are given for the 0-16 kb or 0-2 kb range. Most peaks tend to fall within about 500 bps of either the TSS or the stop codon of a nearby gene. (c) Consensus HRE sequences identified by MEME-Suite in humans and enriched in all *C. elegans* ChIP-seq sequences.

**Extended Data Figure 3.**
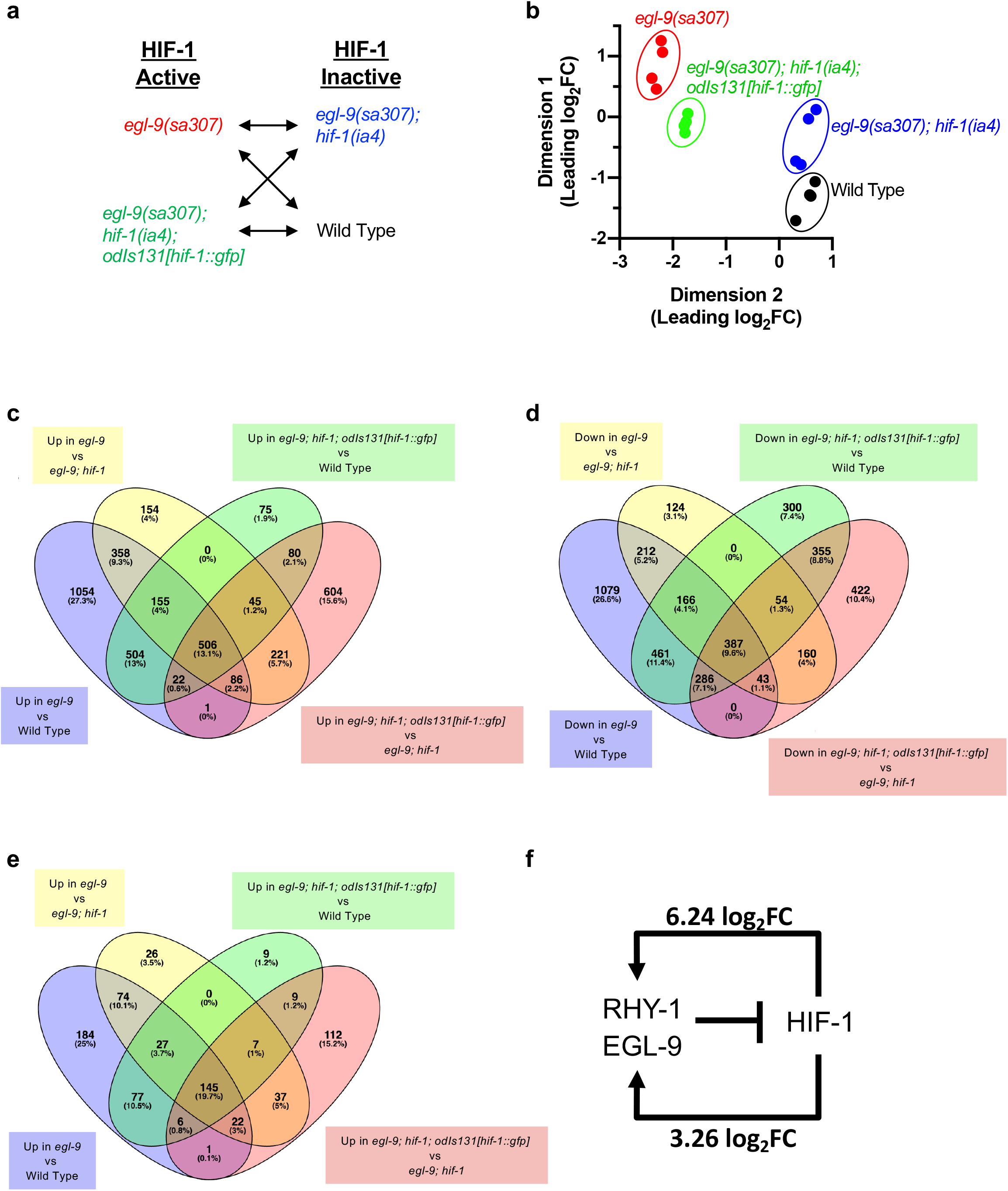
RNA-seq of Several Genotypes Identifies High-confidence Targets. (a) The four different pairwise comparisons of RNA-seq expression data used to identify genes whose expression is changed when HIF-1 is active versus inactive under aerobic conditions. (b) Multidimensional Scaling analysis of RNA-seq gene expression data for four replicates each of four different genotypes. (c-e) Venn diagrams of the number (and percentage of total) of genes with significant differential expression in the indicated pairwise comparison. (c) Genes that are upregulated when HIF-1 is active. (d) Genes with downregulated expression when HIF-1 is active. (e) Direct target genes (i.e., genes with a HIF-1 binding site within 15 kb) that are upregulated when HIF-1 is active. No direct target genes are downregulated when HIF-1 is active. (f) The genes *egl-9* and *rhy-1*, which are negative regulators of HIF-1, are some of the strongest upregulated targets of HIF-1, highlighting the importance of negative feedback regulation in the pathway.

**Extended Data Figure 4.**
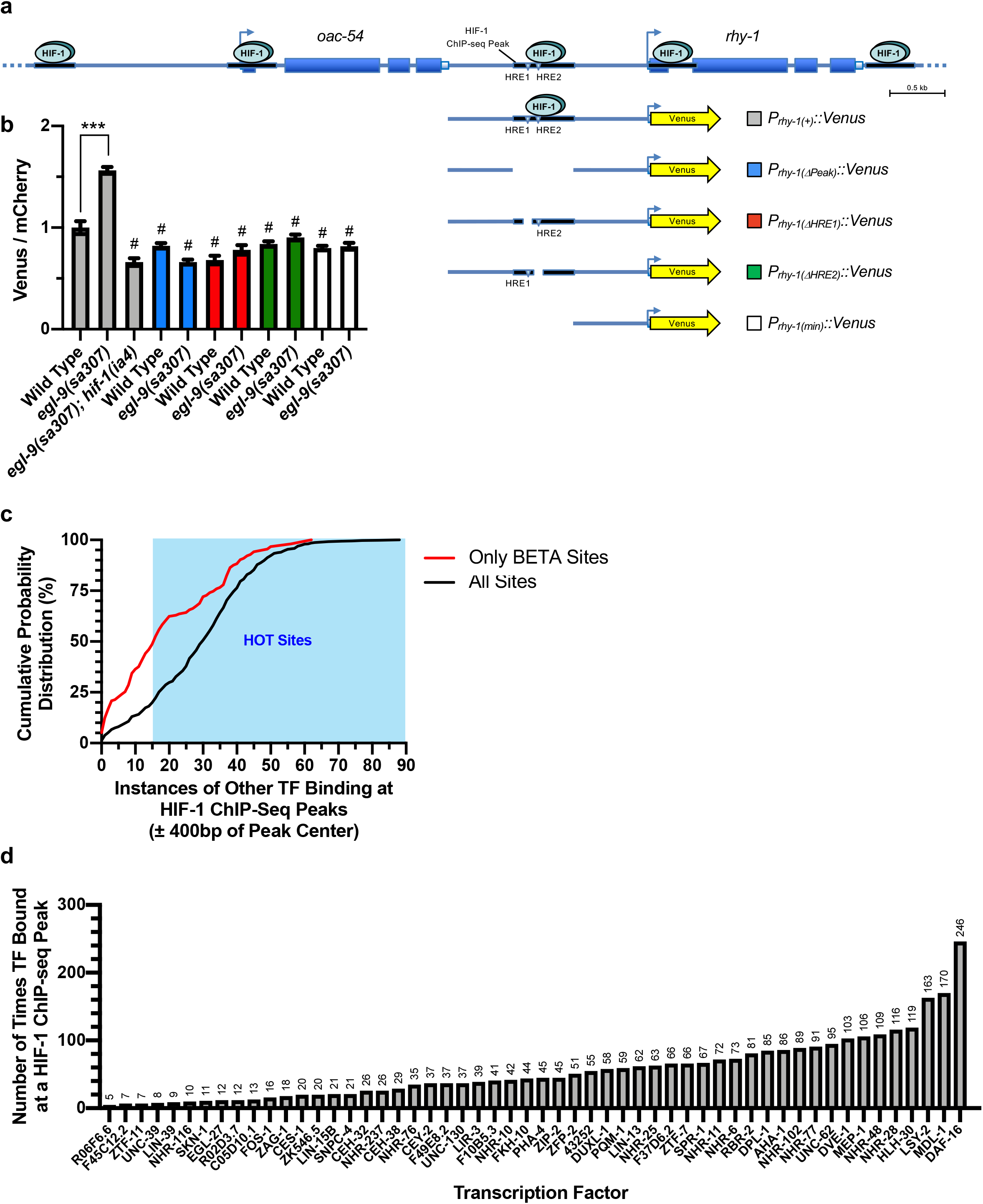
(a) Cartoon of the *rhy-1* locus and the different Venus-based promoters used to examine its regulated expression. Boxes indicate exons. Arrows indicate the TSS. The ovals and black lines beneath them indicate HIF-1 and the HIF-1 binding site identified by ChIP-seq, respectively. The inverted triangles indicate the sites of the HRE motif. The yellow arrow indicates sequences encoding the fluorescent Venus reporter. The *oac-54* gene is nearby this cluster of HIF-1 binding sites and also shows HIF-1-dependent regulation. (b) Graph of Venus/mCherry fluorescence ratios for the indicated genotype. Bar color indicates specific reporter as per panel A. Error bars indicate SEM. ***p<0.001 ANOVA/Dunnett’s Multiple Comparison test compared to wild type. #p<0.001 ANOVA/Dunnett’s Multiple Comparison test compared to *egl-9(sa307)*. (c) Cumulative probability distribution measuring the number of HIF-1 binding sites (ChIP-seq peaks) against the number of other transcription factors known to bind to each site’s region of the genome (within 400 bps of each ChIP-seq peak center). The black line indicates all HIF-1 binding sites, whereas the red line indicates HIF-1 binding sites identified by BETA. Blue shading indicates high occupancy target (HOT) sites determined by modENCODE. (d) Graph of the number of times each indicated transcription factor bound within 400 bps of a HIF-1 binding site (ChIP-seq peak center). Specific number is listed above each bar.

**Extended Data Figure 5.**
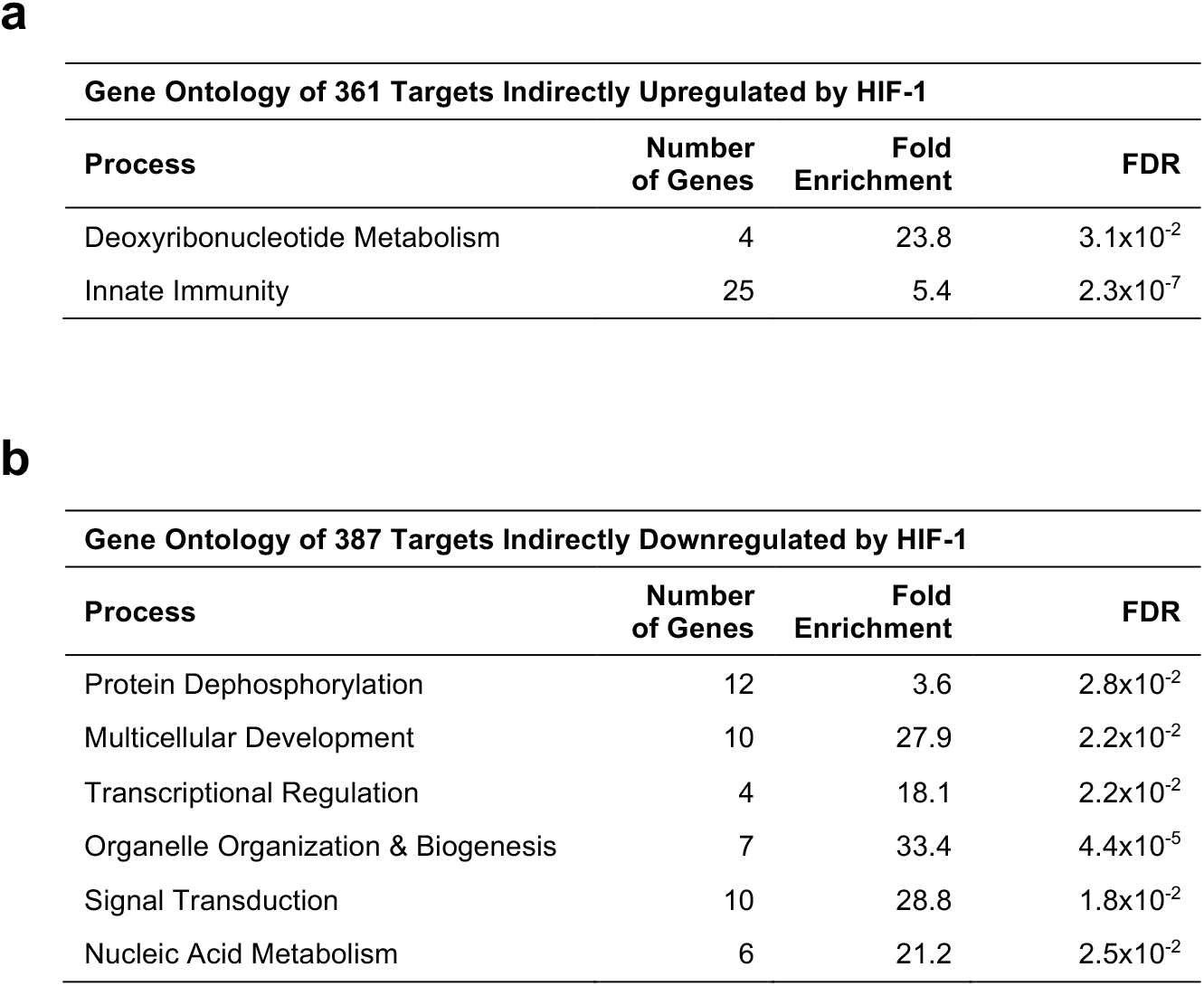
(a,b) Enriched Gene Ontology (GO) classifications for (a) the 361 genes upregulated when HIF-1 is active that are <u>not</u> direct HIF-1 targets, and (b) the 387 genes downregulated when HIF-1 is active. Fold enrichment and FDR for each classification and subclassification is indicated.

**Extended Data Figure 6.**
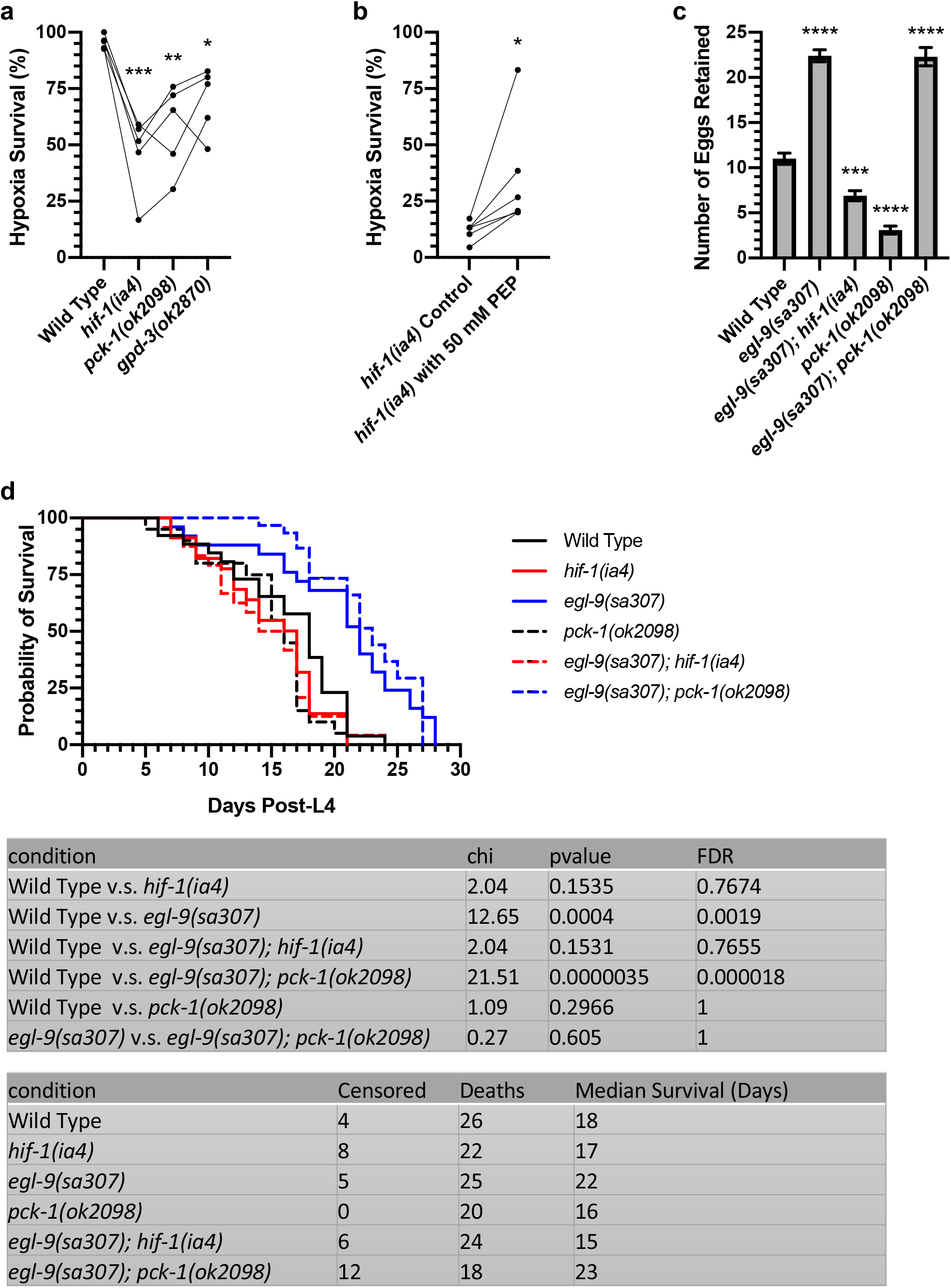
(a,b) Percent of L4-stage animals to survive hypoxia on food (48h 0.1% O2, 25°C, with 24h recovery at 20°C). Because of trial-to-trial variability, survival values for different genotypes within an individual trial are connected by lines. ***p<0.001, **p<0.01, *p<0.05 ANOVA/Dunnett’s multiple comparison test for the average across trials. (c) Average number of unlaid eggs *in utero* per animal for the indicated genotype (n=40-55 animals). ****p<0.0001, ***p<0.001, ANOVA/Dunnett’s multiple comparison test to wild type control. (d) Kaplan-Meier survival curves for animals of the indicated age and for the indicated genotypes. P-values and adjusted FDR values as indicated using the Log-rank test.

## Notes

### Competing Interest Statement

The authors have declared no competing interest.

